# High sensitivity single cell RNA sequencing with split pool barcoding

**DOI:** 10.1101/2022.08.27.505512

**Authors:** Vuong Tran, Efthymia Papalexi, Sarah Schroeder, Grace Kim, Ajay Sapre, Joey Pangallo, Alex Sova, Peter Matulich, Lauren Kenyon, Zeynep Sayar, Ryan Koehler, Daniel Diaz, Archita Gadkari, Kamy Howitz, Maria Nigos, Charles M. Roco, Alexander B. Rosenberg

## Abstract

Single cell RNA sequencing (scRNA-seq) has become a core tool for researchers to understand biology. As scRNA-seq has become more ubiquitous, many applications demand higher scalability and sensitivity. Split-pool combinatorial barcoding makes it possible to scale projects to hundreds of samples and millions of cells, overcoming limitations of previous droplet based technologies. However, there is still a need for increased sensitivity for both droplet and combinatorial barcoding based scRNA-seq technologies. To meet this need, here we introduce an updated combinatorial barcoding method for scRNA-seq with dramatically improved sensitivity. To assess performance, we profile a variety of sample types, including cell lines, human peripheral blood mononuclear cells (PBMCs), mouse brain nuclei, and mouse liver nuclei. When compared to the previously best performing approach, we find up to a 2.6-fold increase in unique transcripts detected per cell and up to a 1.8-fold increase in genes detected per cell. These improvements to transcript and gene detection increase the resolution of the resulting data, making it easier to distinguish cell types and states in heterogeneous samples. Split-pool combinatorial barcoding already enables scaling to millions of cells, the ability to perform scRNA-seq on previously fixed and frozen samples, and access to scRNA-seq without the need to purchase specialized lab equipment. Our hope is that by combining these previous advantages with the dramatic improvements to sensitivity presented here, we will elevate the standards and capabilities of scRNA-seq for the broader community.

## Introduction

The development of microwell^1^, and microfluidic-based^2–4^ single cell RNA sequencing (scRNA-seq) methods has enabled researchers to dissect the cellular composition of tissues and link gene expression signatures to cellular behaviors in health and disease^5–9^. While these initial approaches were extremely informative, they lacked scalability and did not allow for high sample multiplexing, making large projects prohibitively expensive^10^. Combinatorial barcoding based single cell sequencing^11,12^ overcame many of these limitations, making it possible to run larger numbers of samples and cells in a single experiment. With combinatorial barcoding, fixed cells and nuclei go through multiple rounds of split-pool barcoding that ensures molecules within cells receive a unique combination of barcodes, enabling each molecule to be traced back to their cell of origin after sequencing. In addition to increasing throughput, this approach enables a flexible experimental timeline since biological samples can be fixed and stored on different days before being processed in parallel in the same experiment. Contrary to other approaches, combinatorial barcoding does not require any complex microfluidics or other specialized equipment. Over the past few years, this approach has been widely adopted to study developmental processes and disease progression at scale^13–26^.

We previously developed and released the suite of Evercode™ Whole Transcriptome kits (WT Mini, WT, and WT Mega), which dramatically improve the performance and streamline the workflow of the foundational combinatorial barcoding method SPLiT-seq. These kits enable multiplexing of up to 96 samples and profiling of up to 1 million cells in one experiment. To demonstrate these capabilities, we previously generated a 1 million peripheral blood mononuclear cells (PBMCs) dataset and characterized the immune landscape in 12 healthy individuals and 12 patients with type 1 diabetes (Fig.S1).

Even with this level of scalability, improvements to assay sensitivity, leading to increased transcript and gene detection would further sharpen biological resolution. This is especially important in samples and cell types with limited starting RNA content or applications that require detection of specific, low-expression genes. Many of these genes have critical functions that contribute to disease risk^27^ and are needed to accurately characterize cell types^28^.

Here, we introduce the Evercode Whole Transcriptome v2 chemistry which dramatically increases the sensitivity of combinatorial barcoding based scRNA-seq. The v2 chemistry includes improvements to the Evercode Cell Fixation and Nuclei Fixation kits as well as the Evercode Whole Transcriptome kits. Across a variety of sample types, including both cells and nuclei, we find that the new chemistry results in substantial improvements in transcript detection (23% - 162%) and gene detection (8 - 79%) compared to the original Evercode Whole Transcriptome kit (v1 chemistry), while maintaining similarly low doublet rates. The improvement in sensitivity also leads to better clustering, improved assignment of cell types and state, and potential reductions in sequencing costs.

### Validation of Improved Chemistry with a Species Mixing Experiment

To validate the Evercode Whole Transcriptome v2 chemistry, we first performed a species mixing experiment using a mixture of human (HEK293) and mouse (NIH/3T3) cells (Fig. S2A). The two species of cells were first mixed in equal proportions, then fixed, barcoded, and processed into single cell libraries with both the Evercode v1 and v2 chemistries. We found both v1 and v2 chemistries result in low observed doublets, (Fig.S2B, v1: 1.3%; v2: 1.7%) indicating a true doublet rate of less than 4%. Average gene expression was highly correlated across the v1 and v2 chemistries (Fig.S2C, R^2^=0.981 for NIH/3T3 and R^2^= 0.983 for HEK293). Furthermore, joint clustering yielded highly concordant clusters for both chemistries (Fig. S2D), suggesting the chemistry improvements maintain single-cell quality and unbiased gene detection. When we compared sensitivity, we found that the v2 chemistry resulted in a 23% increase in transcript detection and an 8% increase in gene detection at 252,000 reads/cell (Fig.S2E).

### Single Nuclei Profiling of Adult Murine Liver

To test the performance of the v2 chemistry on nuclei, we first chose to test mouse liver, a relatively common yet challenging tissue type. We started with nuclei preparations from a single mouse liver and then divided them into two separate but parallel workflows using the v1 and v2 chemistries (Fig. 2A).

**Figure 1.**
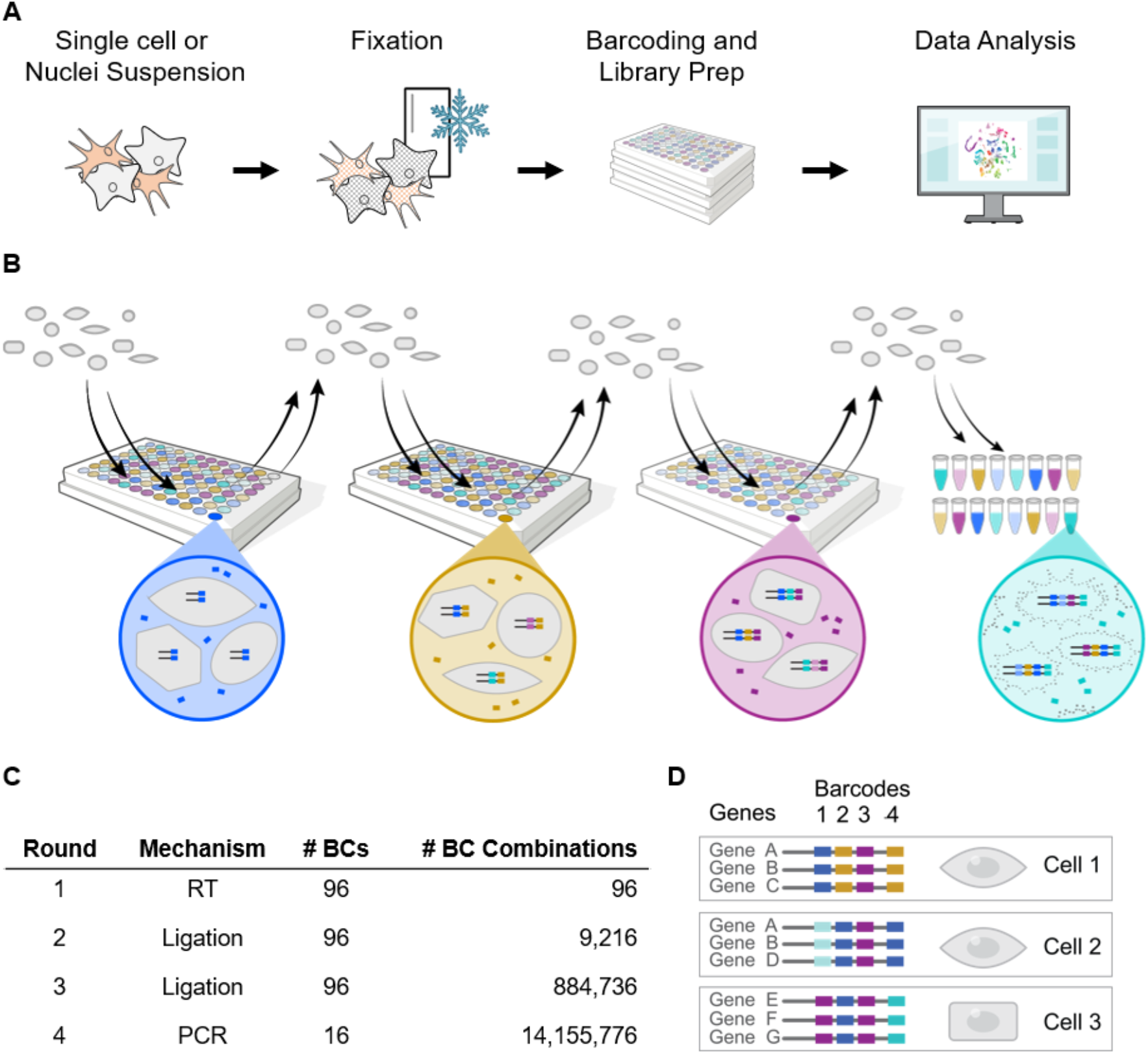
Overview of Evercode Whole Transcriptome workflow. (A) A single cell or nuclei suspension is first fixed, after which fixed cell or nuclei suspensions can be stored before proceeding to barcoding. Barcoding and library preparation is followed by sequencing and data analysis. (B) During the barcoding, fixed cells or nuclei undergo several rounds of split-pool barcoding to uniquely label transcripts according to cell of origin. Cells are then lysed, and barcoded transcripts undergo library preparation, sequencing, and data analysis. (C) With each round of split-pool barcoding, the total number of barcode combinations grows exponentially. The Evercode WT Mega configuration generates over fourteen million barcode combinations, enough to uniquely label one million cells or nuclei with very low doublet rates. (D) Single cell transcriptomes are constructed by grouping transcripts containing the same four barcode combinations.

**Figure 2.**
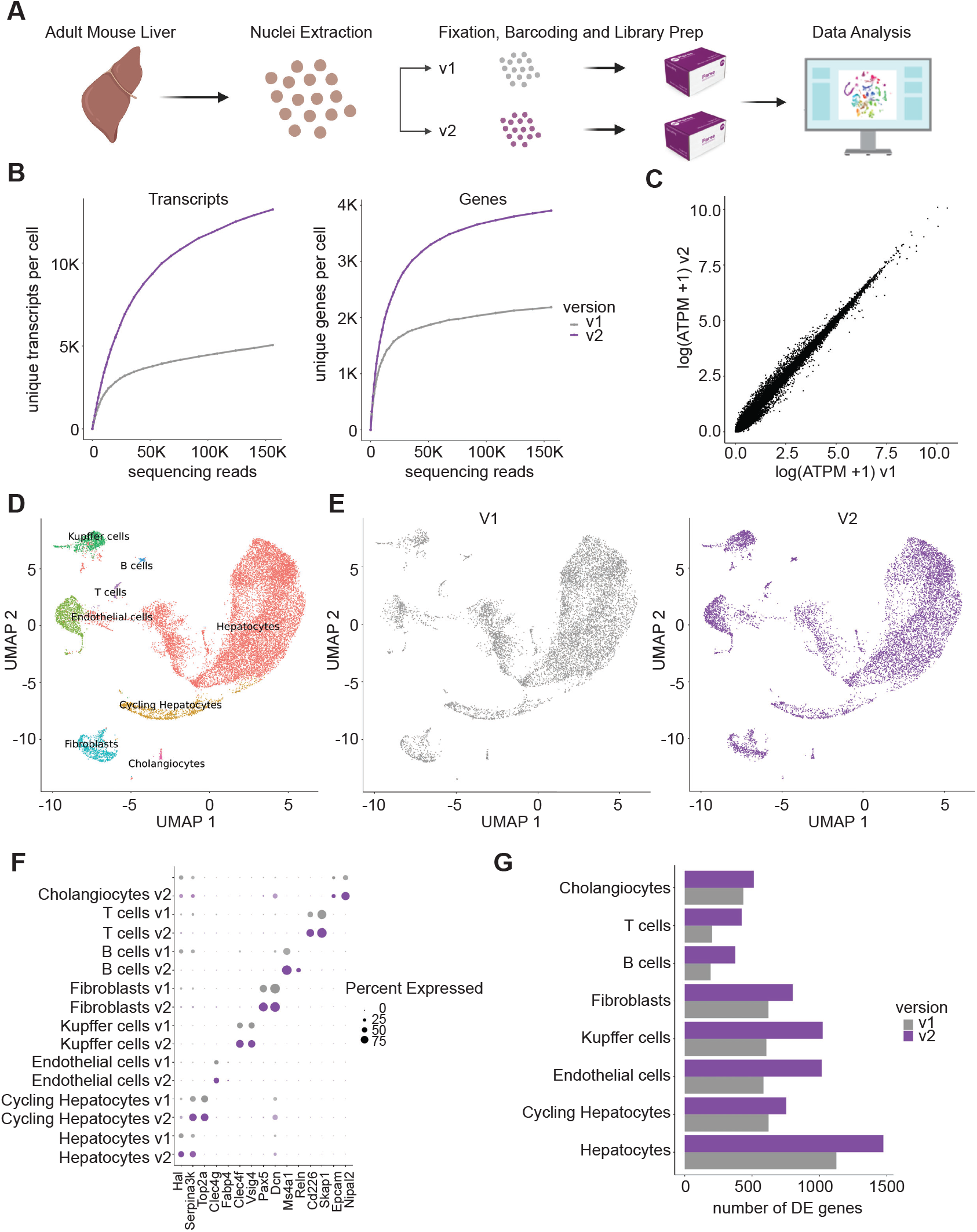
Comparing the performance of Evercode v2 to v1 on nuclei extracted from adult mouse liver. (A) Nuclei were extracted from an adult mouse liver and then fixed and processed in parallel using both the Evercode v1 and v2 chemistries. (B) Number of unique transcripts (left) and unique genes (right) detected at different numbers of raw reads per cell. (C) Correlation of average gene expression (log average transcripts per million plus 1) between the v1 and v2 chemistries. (D) 9,520 nuclei from Evercode v1 and 9,934 nuclei from Evercode v2 were integrated and clustered together and visualized using UMAP (E) Nuclei from v1 and v2 cluster together. (F) Expression of key markers is higher with the v2 chemistry (purple) compared to v1 (gray) (G) v2 chemistry (purple) detects a higher number of differentially expressed genes relative to v1 (gray).

When comparing sensitivity, we found the v2 chemistry resulted in 2.6-fold higher number of transcripts detected per cell (v1: 5,060 transcripts; v2: 13,239 transcripts) and 1.8-fold higher number of genes detected per cell (v1: 2,183 genes; v2: 3,904 genes) when normalized to 156,000 reads/cell sequencing depth (Fig. 2B). Again, we found average gene expression remains highly correlated between v1 and v2 chemistries (Fig. 2C).

After clustering and labeling clusters using documented cell type markers^29^, were able to identify all major cell types for both the v1 and v2 chemistry. (Fig. 2D) The proportions of cell types were also similar between the chemistries and in line with previously reported cell type proportions (Figs. 2E, S3A). Approximately three quarters of identified cells were hepatocytes with smaller populations corresponding to cycling hepatocytes, endothelial cells, Kupffer cells, fibroblasts, cholangiocytes, T cells, and B cells. Key marker genes across different cell types were more highly and uniformly detected using the v2 chemistry (Figs. 2F, S3B) and the v2 chemistry consistently detected more differentially expressed genes (Fig. 2G).

### Sensitive and Accurate Profiling of PBMCs

We then performed a comparison of the v2 and v1 chemistries on peripheral blood mononuclear cells (PBMCs) (Fig. 3A). PBMCs are typically small and thus contain less mRNA content, which often makes it challenging to detect high numbers of genes per cell. In addition to measuring the sensitivity of the v2 and v1 chemistries, we assessed consistency of gene detection across different samples and the accuracy of recovered cell type proportions. We obtained PBMC samples from four different donors and processed each PBMC sample in parallel with both the v1 and v2 chemistries. We sequenced and analyzed 2,500-4,000 cells from each PBMC sample for each chemistry, resulting in the analysis of ~12,000 total cells across donors for each chemistry.

**Figure 3.**
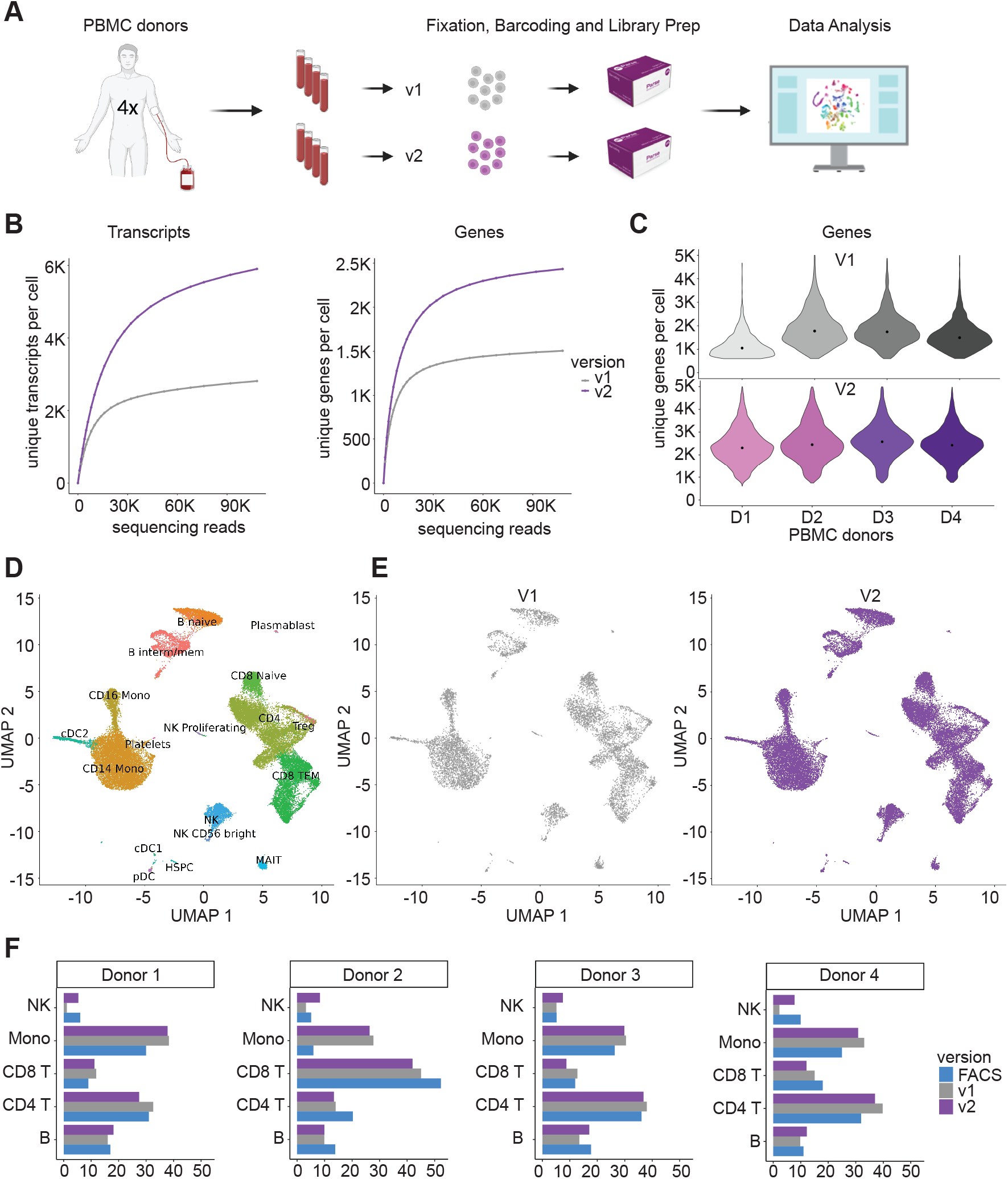
Profiling human peripheral blood mononuclear cells from different donors with Evercode v2. (A) PBMCs from four different donors were fixed and processed in parallel using both the Evercode v1 and v2 chemistries. (B) Number of unique transcripts (left) and unique genes (right) detected at different numbers of raw reads per cell. (C) Violin plot showing the number of detected genes per PBMC donor (D1-D4) for Evercode v1 (top) and Evercode v2 (bottom). The dot (black) denotes the median genes detected for each donor and demonstrates Evercode v2 increases both the number of detected genes and reduces sample to sample variability. (D) 10,134 cells from Evercode v1 and 15,201 cells from Evercode v2 were integrated and clustered together. (E) Cells prepared using either the v1 or v2 chemistry cluster similarly. (G) Both the v1 and v2 chemistries result in concordant proportions of cell types as compared to flow cytometry.

When we compared the sensitivity between v1 and v2, we found that, on average, v2 chemistry detected 2.1-fold more transcripts per cell (v1: 2,812 transcripts; v2: 5,903 transcripts) and 1.6-fold more genes per cell (v1: 1,506 genes; v2: 2,434 genes) (Fig.3B). The improvement in sensitivity was also consistent across different cell types (Fig. S5B). We generally found that key immune marker genes were more highly expressed as well, making it easier to identify and label different populations of cells (Fig. S5C). We also found more consistent gene detection per cell across the different PBMC samples for the v2 chemistry compared to the v1 chemistry. At 76,000 reads per cell, gene detection for the v1 chemistry varied between 935 to 1,682 (Δ=747) while gene detection was between 2,233 to 2,493 (Δ=260) for the v2 chemistry (Figs. 3C, S5A). This observation suggests that the v2 chemistry may mitigate technical variability between samples.

We next measured the concordance between cell type proportions identified by single cell sequencing for both the v1 and v2 chemistries and independent measurements from flow cytometry. To do this, we jointly clustered both single cell datasets using Seurat and labeled each cluster with Azimuth using previously published reference datasets^30^ (Figs. 3D, 3E). We then calculated the proportions of each major cell type and compared the results to data collected using flow cytometry. Across all cell types, we found high concordance both between the v1 and v2 chemistries and flow cytometry, indicating unbiased recovery of PBMC cell types for both chemistries (Fig. 3F).

### Single Nuclei Profiling of the Developing Murine Brain

We tested the performance of the Evercode Whole Transcriptome v2 chemistry on single nuclei sequencing of embryonic mouse brains. We extracted nuclei from two different flash frozen E18 mouse brains which were subsequently fixed and processed using both the v1 and v2 chemistries in parallel (Fig.4A).

**Figure 4.**
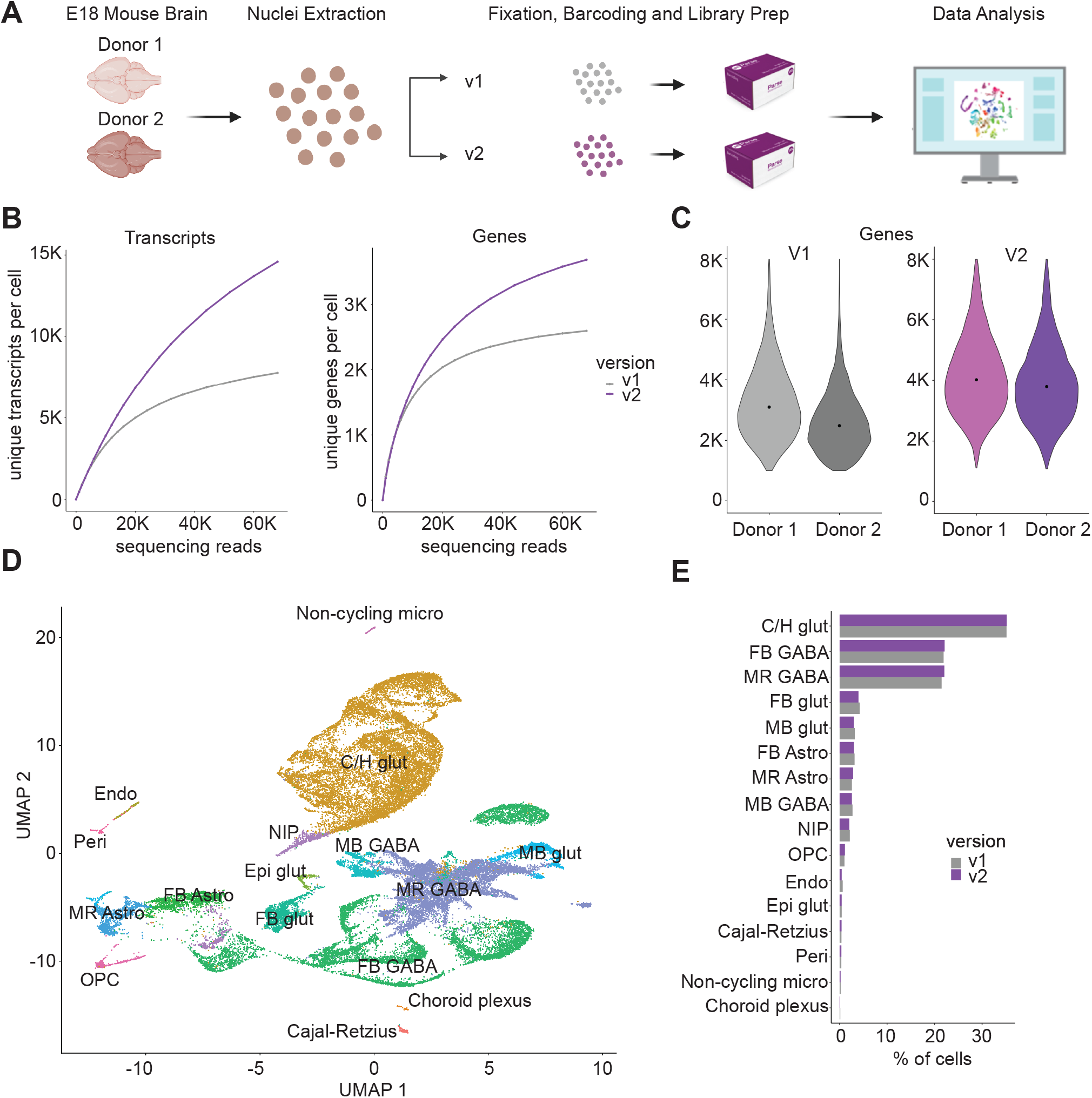
snRNA-seq profiling of two E18 mouse brains with Evercode v2. (A) Nuclei were extracted from two different E18 mouse brains and then processed in parallel using both Evercode v1 and v2 chemistries (B) Number of unique transcripts (left) and unique genes (right) detected at different numbers of raw reads per cell. (C) Violin plot showing the number of detected genes per mouse brain donor for Evercode v1 (left) and Evercode v2 (right). The dot (black) denotes the median genes detected per donor. (D) 24,011 cells from Evercode v1 and 18,524 cells from Evercode v2 were integrated and clustered together. (E) Cells from different donors cluster similarly as do cells prepared using either the v1 or v2 chemistry. (F) Cell type proportions are the same between the Evercode v1 (gray) and v2 (purple) chemistry.

As with the other samples profiled, we found the Evercode v2 chemistry resulted in a dramatic increase in both transcripts per cell (Fig.4B, v1: 7,768 transcripts; v2: 14,588 transcripts) and genes per cell (Fig. 4B, v1: 2,595 genes; v2: 3,686 genes). The v1 chemistry had larger variability in gene detection than the v2 chemistry between the two brain samples (Figs. 4C, S6A), v1: 2,345 and 2,846 genes, v2: 3,768 and 3,605 genes), indicating that the v2 chemistry results in increased consistency across different samples.

We jointly clustered ~40,000 nuclei split equally between v1 and v2. The clusters were labeled with Seurat, using the developing mouse brain atlas from La Manno *et al*^31^ (Fig. 4D). When comparing cells from the two different mouse brains, we found similar clustering (Fig. S6E). A comparison of cell types identified between the v1 and v2 chemistries also showed highly similar proportions across all cell types, suggesting that the v2 chemistry maintains unbiased detection across cell types (Fig. 4E).

## Discussion

As single cell sequencing has matured, advances such as combinatorial barcoding have enabled larger scale experiments^11,12^. This has made it possible to design experiments with greater numbers of biological samples and replicates while simultaneously increasing the number of cells per sample. Including replicates is especially important to ensure reproducibility across experiments^32^ and increasing the number of cells per sample enables profiling of rare cell subpopulations^8^. While increasing the number of cells profiled can be powerful, previous work has also highlighted the importance of increasing the sensitivity of high-throughput single cell sequencing^33,34^.

To address this problem, we have introduced the Evercode Whole Transcriptome v2 chemistry, which dramatically improves the sensitivity of the Evercode single cell platform. We found improved transcript and gene detection across cell lines, human PBMCs, and nuclei extracted from both mouse liver and mouse brain tissue. We also found that the Evecode Whole Transcriptome v2 chemistry improves the consistency of single cell results across different biological samples in the same experiment.

The increased sensitivity of the v2 chemistry adds to the existing advantages of the Evercode single cell platform. Leveraging the high multiplexing and exponential scalability of combinatorial barcoding makes it possible to scale projects to profile millions of cells and hundreds of samples. There is also no need to purchase a custom instrument, increasing accessibility. The ability to fix and store samples on different days and then barcode at a later time offers further flexibility around experimental design. Combining these advantages with dramatically improved sensitivity makes Evercode a powerful tool for researchers using single cell sequencing.

## Materials and Methods

### Preparation of Cultured Cells for Single Cell Sequencing

Adherent HEK293 cell lines were grown in 10 mLs of DMEM (ATCC 302002) containing 10% FBS (Gibco 16000044) and 1% Pen/Strep (Gibco 15070063). Adherent NIH/3T3 were grown in 10 mLs DMEM with 10% BCS (Sigma 12133C) and 1% Pen/Strep. Plates were harvested at 85% confluence using TrypLE Express (Thermo Fisher 12605010).

HEK293 and NIH3T3 cells were counted using a hemocytometer and combined in an even 50:50 ratio. Combined cells were centrifuged at 200 xg for 10 min, the supernatant was discarded, and cells were resuspended in 750 μL of Cell Prefixation Buffer. Samples were then fixed using the Parse Biosciences Cell Fixation (v1 or v2) kit and stored at −80°C until the start of barcoding and library prep with the Evercode Whole Transcriptome (v1 or v2) kit.

### Preparation of Human Peripheral Blood Mononuclear Cells (PBMCs) for Single Cell Sequencing

Cryopreserved PBMC samples were obtained from Bloodworks Northwest. Cells were thawed in a water bath set to 37°C, then revived by dropwise addition of 50 mLs of warm media (RPMI (Gibco 11875093) +10% FBS (Gibco 16000044)): first by adding 1mL at the rate of 1 drop/5 seconds, then 1 mL/5 seconds in increments of 2 mL, 4 mL, 8 mL, 16 mL, and 18 mL (with one minute incubation time between each round of media addition). The cells were gently swirled during media addition. Cells were centrifuged at 200 x g for 10 minutes, and the supernatant discarded. Cells were then resuspended in 10 mLs of cold media (RPMI +10% FBS, Gibco 11875093, Gibco 16000044) and counted via automated cell counter using trypan blue dye. Cells were centrifuged again at 200 x g for 10 minutes. Supernatant was discarded and pellets were resuspended in 750 μL of Cell Prefixation Buffer. Samples were then fixed using the Parse Biosciences Cell Fixation (v1 or v2) kit and stored at −80°C until the start of barcoding and library prep with the Evercode Whole Transcriptome (v1 or v2) kit.

### Preparation of Murine Brain Nuclei for Single Cell Sequencing

Flash frozen E18 brains from C57/Bl6 mice were obtained from TransNetYX Inc (Springfield, IL) and stored in liquid nitrogen. Prior to flash freezing the brain, the meninges was removed and the whole brain cut down the longitudinal fissure in the sagittal plane resulting in a left and right hemisphere for each donor. Prior to performing nuclei fixation for the Parse Biosciences fixation protocol, nuclei from the left hemisphere of donor 1 and the right hemisphere of donor 2 were isolated using a standard douncing method. Prior to and during nuclei isolation, all tools and buffers were kept chilled on ice. Preparation for nuclei isolation buffer 1 (NIM1) includes reagents at the following final concentration: 250mM Sucrose (Sigma S0389), 25mM KCl (Invitrogen AM9640G), 5mM MgCl2 (Invitrogen AM9530g), 10mM Tris pH8 (Invitrogen AM9856), and nuclease free water. Nuclei isolation buffer 2 (NIM2) derived from NIM1 buffer by adding the following to the final concentrations: 1uM DTT (Invitrogen P2325), 0.4U/μl Enzymatic RNase-In (Qiagen Y9240), 0.2U/μl Superase-In (Invitrogen AM2696), and 0.1% Triton X-100 (Sigma T8787).

A swinging bucket centrifuge was pre-cooled to 4°C prior to starting nuclei isolation. Frozen brain sample was placed in 700μL of cooled NIM2 buffer, transferred to a dounce, and homogenized with 10 strokes using the loose pestle and 10 strokes using the tight pestle. The nuclei homogenate was transferred and filtered through a 40μm filter into a 15ml conical tube and nuclei were counted using a hemocytometer. The sample was partitioned to ensure there were less than 5 million nuclei in each aliquot and centrifuged at 200 x g for 10 min at 4°C. Pellets were resuspended in Parse Nuclei Buffer containing 0.75% BSA and then fixed using the Parse Biosciences Nuclei Fixation (v1 or v2) kit and stored at −80°C until the start of barcoding and library prep with the Evercode Whole Transcriptome (v1 or v2) kit.

### Preparation of Murine Liver Nuclei for Single Cell Sequencing

Flash frozen adult liver from C57/Bl6 mouse was obtained from TransNetYX Inc (Springfield, IL) and stored in liquid nitrogen. Prior to flash freezing, the liver was segmented into equal sections. As previously described for brain nuclei isolation, NIM2 buffer was made fresh before nuclei isolation. Tissue was removed from liquid nitrogen storage and thawed on ice in 500μL of cooled NIM2 buffer for 5 minutes. A swinging bucket centrifuge was pre-cooled at 4°C prior to starting nuclei isolation. Sterile and chilled scissors were used to mince the tissue while still in the NIM2 buffer. Minced liver is transferred to a dounce, NIM2 buffer was added to a total of 700μL, and homogenized with 10 strokes using the loose pestle and 15 strokes using the tight pestle. The nuclei homogenate was filtered through a 70 μm filter into a 15ml conical tube and spun down at 500 x g for 3 minutes. The pellet was resuspended in 1 mL of NIM2 buffer and filtered through a 40μm filter. The sample was counted to ensure less than 5 million nuclei in each aliquot. The samples were centrifuged at 200 x g for 10 min at 4°C and were resuspended in Parse Nuclei Buffer containing BSA. Nuclei were then fixed using the Parse Biosciences Nuclei Fixation (v1 or v2) kit and stored at −80°C until the start of barcoding and library prep with the Evercode Whole Transcriptome (v1 or v2) kit.

### Barcoding and Library Preparation

In advance of cell barcoding and library preparation, fixed samples were removed from −80°C, thawed in a water bath set to 37°C, placed on ice, and counted. Samples were barcoded and single cell sequencing libraries were then prepared using the Evercode Whole Transcriptome (v1 or v2) kit.

### Parse v1 and v2 sequencing and data processing

For the mixed species experiment, libraries were sequenced on a Next-seq 550 Illumina instrument using two high output 150 cycle kits. For the mouse liver nuclei experiment, libraries were also sequenced on a Next-seq 550 Illumina instrument using two high output 150 cycle kits. E18 mouse brain nuclei and PBMC libraries were sequenced on a Novaseq 6000 Instrument using the S4 kit. Sequencing reads from the mRNA libraries were mapped to human (hg38) or mouse (mm10) genomes using the parse biosciences pipeline (split-pipe v0.9.6p) to generate cell by gene counts matrices. To ensure v1 and v2 chemistry comparisons were done fairly, we downsampled all libraries to normalize everything to the same number of mean reads per cell. Count matrices were then used as input for the Seurat R package^30^ to perform all downstream analyses.

### QC and data processing and clustering for the species mixing experiment

Cells with low quality metrics such as high mitochondrial gene content (> 5%) and low number of genes detected (<600) were removed. Cells with transcripts from both mm10 and hg38 were removed as doublets. RNA counts were log normalized using the standard Seurat workflow. To visualize cells based on an unsupervised transcriptomic analysis, we first ran PCA using 2,000 variable genes. For this experiment the first 10 components were used as input for UMAP visualization in two dimensions.

### QC and data processing and clustering for the adult mouse liver nuclei experiment

Cells with low quality metrics such as high mitochondrial gene content (> 15%) and low number of genes detected (<600) were removed. In an attempt to also remove doublets we filtered out cells with more than 7,500 genes detected. RNA counts were log normalized using the standard Seurat workflow. For this experiment the first 10 components were used as input for UMAP visualization in two dimensions.

To visualize cells based on an unsupervised transcriptomic analysis, we first ran PCA using 2,000 variable genes. The first 10 components were used as input for UMAP visualization in two dimensions.

### QC and data processing and clustering for the PBMC experiment

Cells with low quality metrics such as high mitochondrial gene content (> 5%) and low number of genes detected (<600) were removed. In an attempt to also remove doublets we filtered out cells with more than 5,000 genes detected. RNA counts were log normalized using the standard Seurat workflow. To jointly analyze all 4 PBMC donors, we used the Seurat fast integration method (rPCA). After integration, we obtained level 2 cell type annotations using the Azimuth app. During this process, a cell was assigned the Azimuth annotation if the prediction score was higher than 0.5. Additionally, we leveraged Leiden cluster assignments to find the most frequent assignment for each cluster and give cells within that cluster that assignment. Finally, if the most frequent assignment within a cluster did not represent more than 25% of cells in that cluster, we removed the cluster assignments.

To cluster and visualize cells based on their transcriptome, we first scaled the integration assay and then ran PCA using 2,000 variable genes. Next, we used the first 40 PCs to build an SNN graph using the FindNeighbors function in Seurat clustered the data using FindClusters with the resolution parameter set to 0.6. Finally, the first 40 components were used as input for UMAP visualization in two dimensions.

### QC and data processing and clustering for the E18 mouse brain nuclei experiment

Cells with low quality metrics such as high mitochondrial gene content (> 3%) and low number of genes detected (<1,000) were removed. In an attempt to also remove doublets we filtered out cells with more than 8,000 genes detected. RNA counts were log normalized using the standard Seurat workflow. To cluster and visualize the mouse brain nuclei based on their transcriptome, we ran PCA using 2,000 variable genes. Next, the first 50 components were used as input for UMAP visualization in two dimensions. While there were 2 mouse brain donors, we observed no donor specific clustering and decided to proceed without donor integration.

To identify all cell types present in our data we performed reference-based mapping and annotation using the mouse brain nuclei from previous literature as a reference^31,35^. We retained annotations with a prediction score higher than 0.5. Furthermore, we leveraged leiden cluster assignments to find the most frequent assignment for each cluster and give cells within that cluster that assignment. Finally, if the most frequent assignment within a cluster did not represent more than 25% of cells in that cluster, we removed the cluster assignments.

## Data availability

The datasets demonstrated in this study will be made available for viewing and download at www.parsebiosciences.com/datasets.

## Supplemental Figures

**Figure S1.**
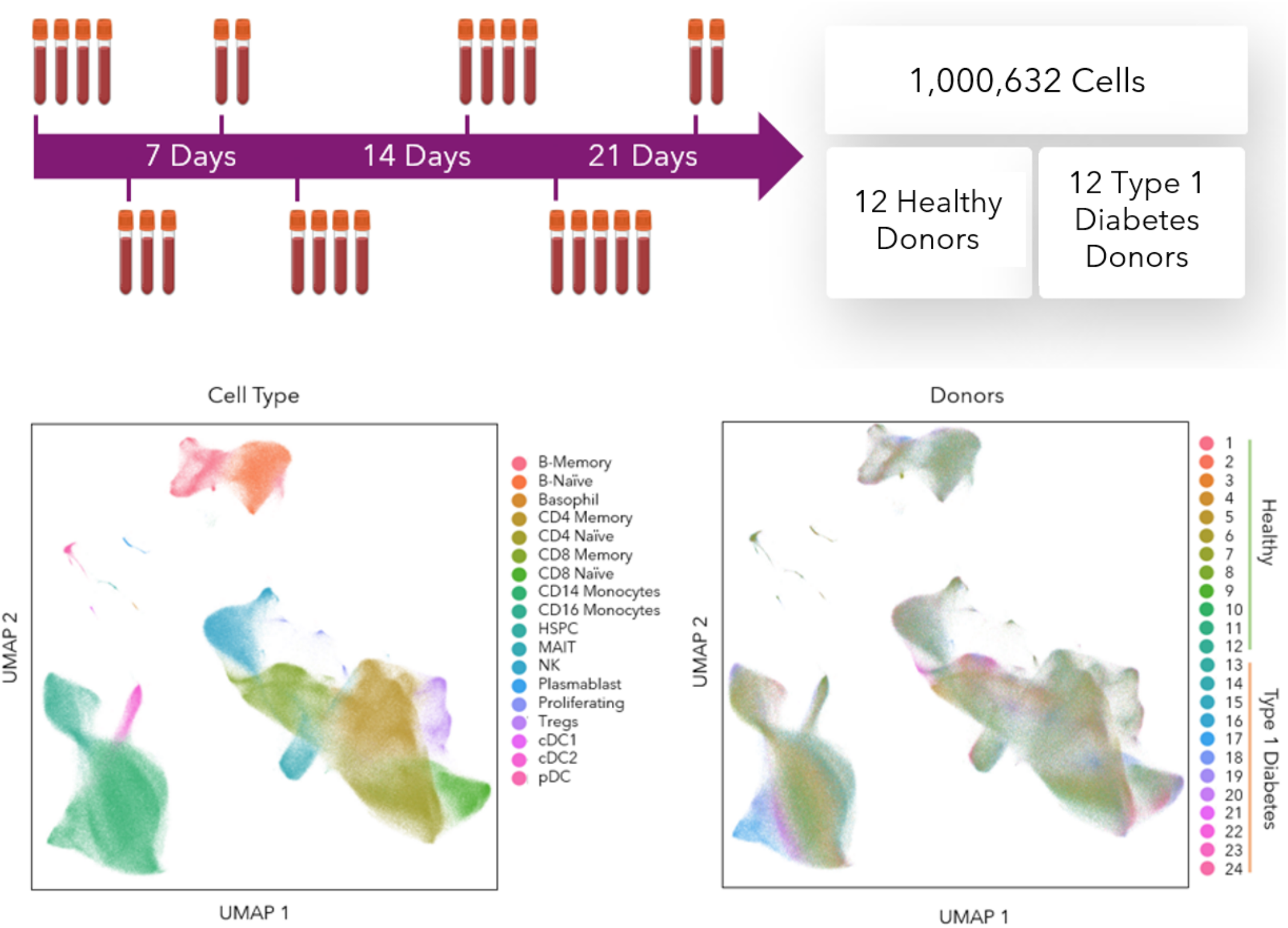
Demonstration of profiling one million cells in a single experiment using combinatorial barcoding. Twenty-four PBMC samples from both healthy and type 1 diabetic donors were collected over the course of three weeks. Using the v1 chemistry, samples were fixed at the time of collection and were subsequently stored. After all samples were collected, an Evercode Whole Transcriptome Mega v1 kit was used to profile over 1 million cells from the 24 samples. UMAP plots reveal cell types that were detected (left) and allowed us to overlay donor specific information, further split by healthy vs type 1 diabetes (T1D) donors (right).

**Figure S2.**
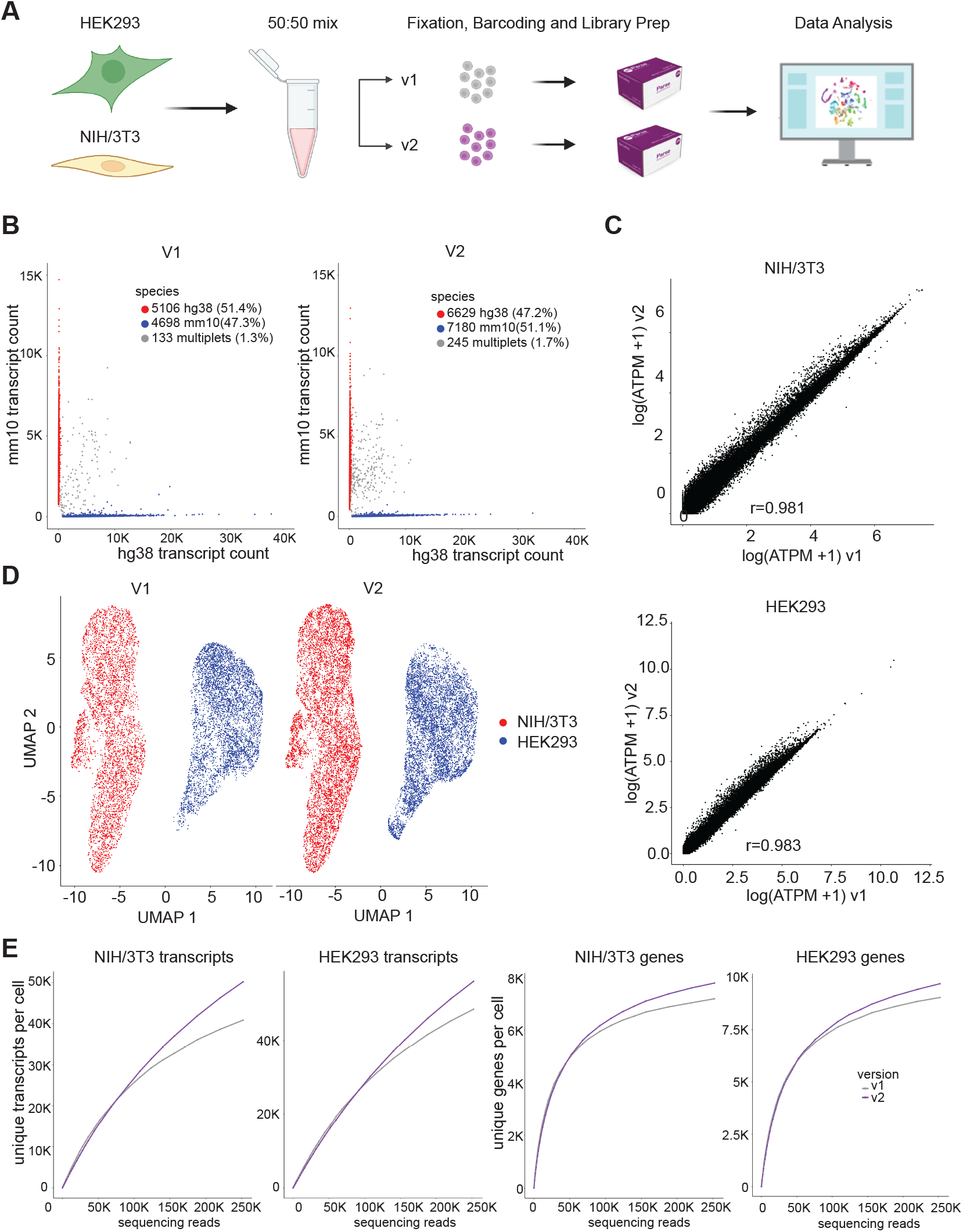
Validation of the Evercode v2 chemistry with a species-mixing experiment. (A) An equal mixture of human (HEK293) and mouse (NIH/3T3) cells were fixed, barcoded, and processed into single cell libraries using both the v1 and v2 chemistries. (B) Species-mixing experiment showing the number of unique human transcripts per cell (x-axis) and unique mouse transcripts per cell (y-axis). Observed doublets are <2% for both the v1 and v2 chemistries, indicating an actual doublet rate of <4% (with unobserved mouse-mouse and human-human doublets accounted for). (C) Correlation of average gene expression (log average transcripts per million) between the v1 and v2 chemistries for both mouse genes (left) and human genes (right). (D) UMAP clustering of species-mixing experiment showing cells from v1 (left) and v2 (right) chemistries co-cluster. (E) Transcripts (left) and genes (right) detected per cell for HEK293 and NIH/3T3 for both the v1 and v2 chemistries.

**Figure S3.**
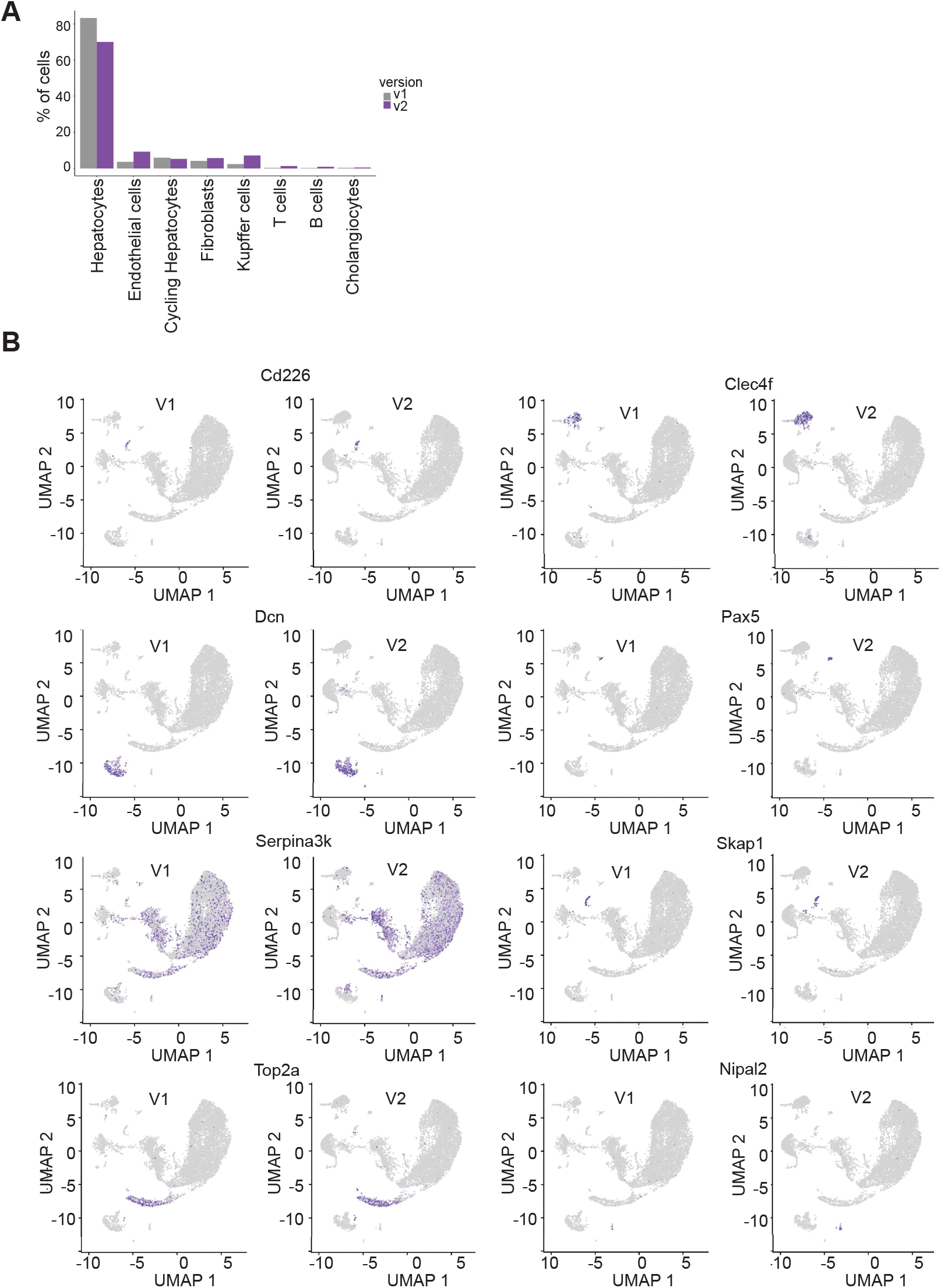
Analysis of cell types and expression levels in liver dataset. (a) Proportion of cell types for v1 and v2 chemistries. (b) UMAP plots showing expression of representative genes for the v1 and v2 chemistries in the liver dataset.

**Figure S4.**
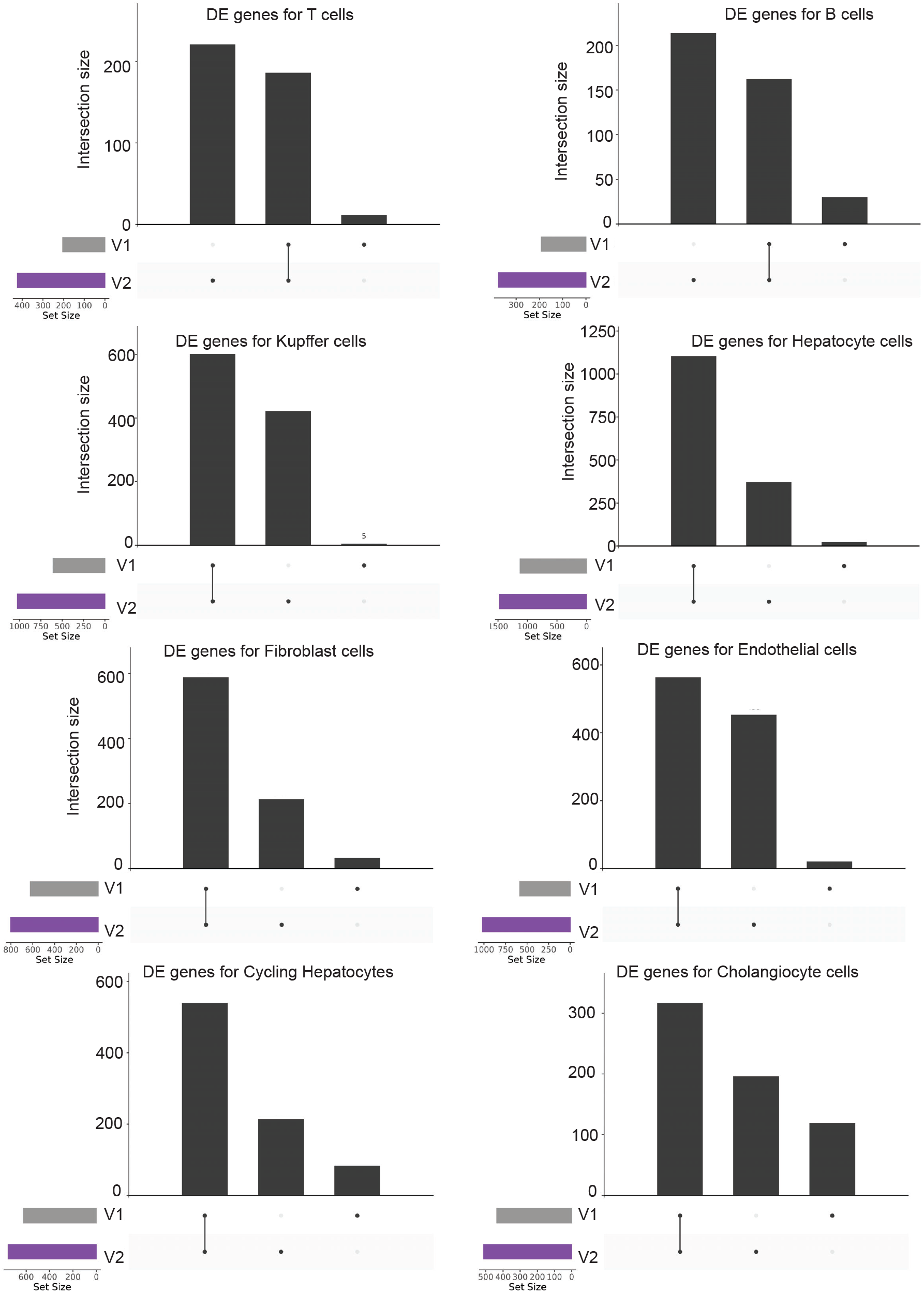
Intersection between v1 and v2 differentially expressed genes across cell types.

**Figure S5.**
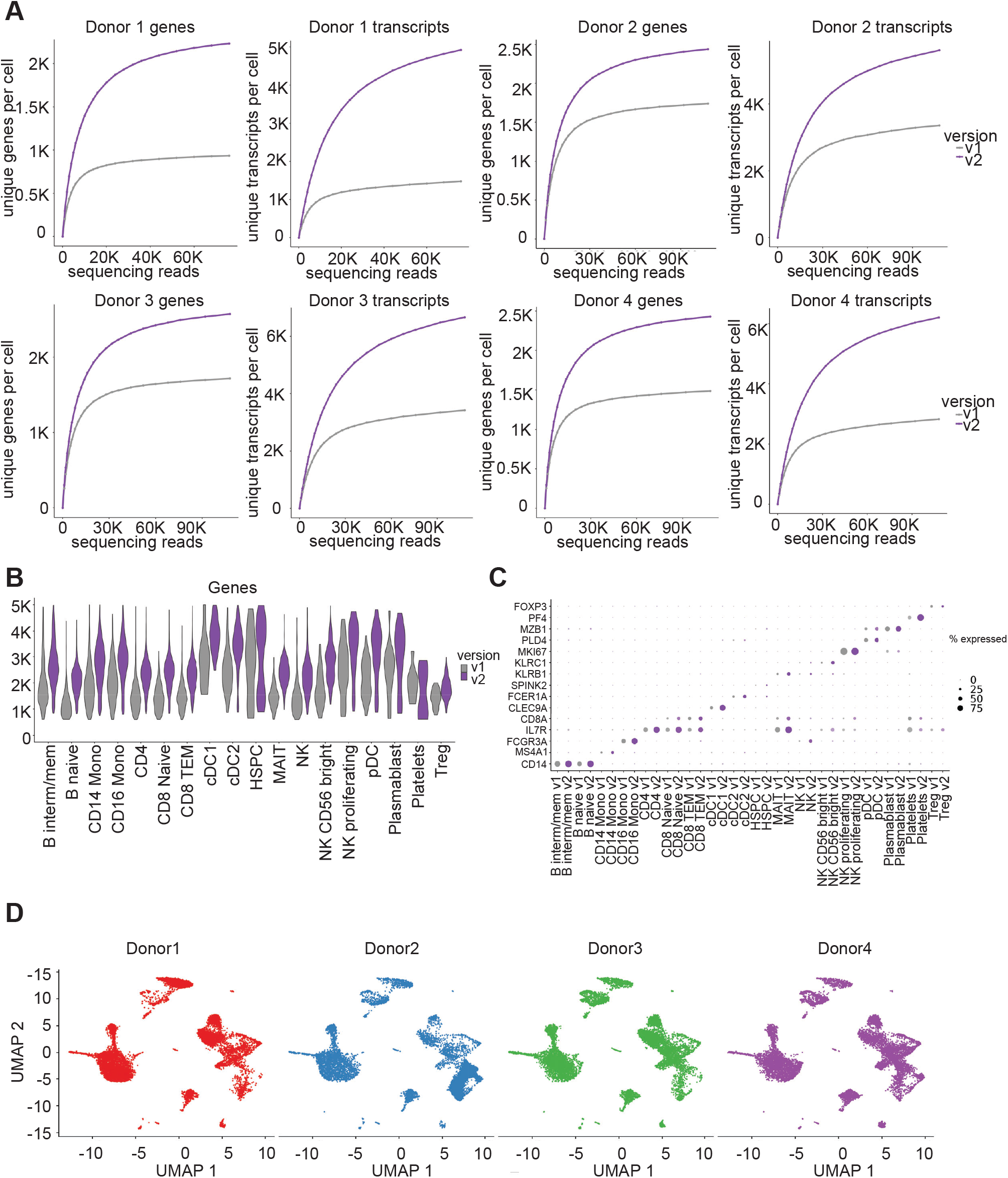
Transcript and gene detection per cell across different PBMC donors. (a) Number of unique transcripts (left) and unique genes (right) detected at different numbers of raw reads per cell across four individual donors. (b) Gene detection of v2 chemistry is higher for all cell types. (c) Expression of key markers is higher with the v2 chemistry (purple) compared to v1 (gray). (d) UMAP clustering of each donors (1-4)

**Figure S6.**
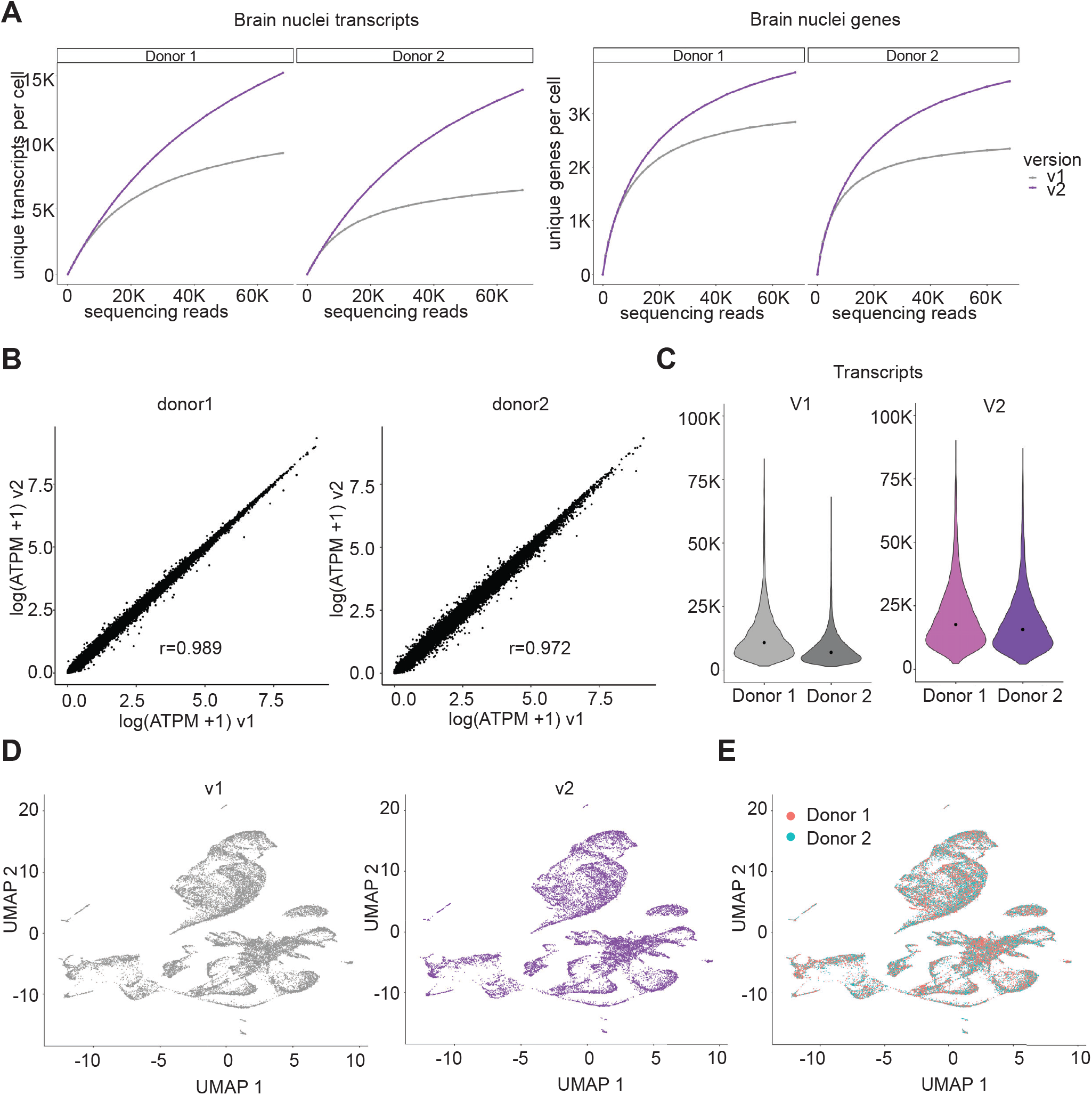
Gene expression across different mouse brain samples. (A) Number of unique transcripts and unique genes detected at different numbers of raw reads per cell across two different brain samples for both v1 and v2 chemistries. (B) Comparison of average gene expression (log average transcripts per million) between the v1 and v2 chemistries. (C) Violin plots of transcript detection for both mouse brains. (D) UMAP clustering shows cells from v1 (left) and v2 (right) chemistries co-cluster. (E) UMAP clustering for Donors 1 and 2 are similar.

## Notes

### Competing Interest Statement

V.T, E.P, S.S, G.K, A.S, J.P, A.S, L.K, Z.S, R.K, D.D, A.G, K.H, M.N, C.R, and A.R are currently employed by Parse Biosciences, some of whom are inventors on granted patents or patent applications pertaining to split pool combinatorial barcoding.

